# A longitudinal study of perceived stress and cortisol responses in an undergraduate student population from India

**DOI:** 10.1101/867184

**Authors:** Anuradha Batabyal, Anindita Bhattacharya, Maria Thaker, Shomen Mukherjee

## Abstract

Young adults entering into college experience immense shifts in both personal and professional environments and this may result in some of them experiencing a lot of stress and difficulty in coping with their new surroundings. Such potentially stressful events may trigger multiple psychological as well as physiological effects. The current study investigated multiple psychological parameters such as PSS14 (Perceived Stress Scale), K10 (distress scale) and positive mood measures, along with salivary cortisol levels, in a repeated measures longitudinal study of first year students (~ 19 years of age) enrolled at a residential college in India. Six salivary cortisol samples were collected over a one-year period from 20 students. On each sampling day, a questionnaire designed to evaluate (K10, PSS14 and Mood) psychological parameters was also administered.

Overall, men showed a significantly lower level of salivary cortisol compared to women. Men also showed a decrease in perceived stress (PSS14) and distress (K10) with time. However, women reported similar perceived stress and distress levels all year round. Academic stress was reported by the students to be the most important stressor, whereas financial stress was reported the least number of times by all participants. Our results suggest that men seem to have a better capability to adjust to the new environment of a residential program with time. In contrast, women show an elevation in salivary cortisol at the end of the semester (the final assessment stage) in spite of a continuous assessment curricular design. This study not only provides an important glimpse into the sex differences in stress response in the first one year of joining an undergraduate program, but it also provides a valuable longitudinal dataset from the Indian undergraduate student cohort which is lacking in literature.

## Introduction

Stress is any event that poses a threat or challenge to physical and mental wellbeing of an individual (Lazarus and Folkman, 1984). Emerging adults starting college, face stressful events such as coping with a new academic environment, relational responsibility, future financial security and, searching for their own identity among others (Kadison and DeGeronimo, 2004). Late adolescence is a critical age where stressors can affect the physiology and psychology of individuals and risk development of mental health issues in the future (Tennant, 2002). Early life stressors can lead to the onset of anxiety symptoms (Breslau et al., 1997), depression (Brown et al., 1996), schizophrenia (Patel et al., 2007) and even suicidal tendencies (Wilcox et al., 2010). The major physiological reaction mechanism by which individuals cope with any stressor is the activation of Hypothalamo-Pituitary-Adrenal (HPA) axis. Secretion of glucocorticoid hormones especially cortisol from an activated HPA axis mediates a suite of physiological responses that has immediate adaptive function to reduce the impact of the stressor. But prolonged and repeated encounter with stressors leads to dysregulation of the HPA axis, causing detrimental effects on multiple organs and systems (Bollini et al., 2004; López et al., 1999; Sapolsky, 1996; Tsigos and Chrousos, 2002). Chronic stress can result in hyper secretion of cortisol thus downregulating receptor numbers which results in lower negative feedback to the hypothalamus and in one extreme can lead to exaggerated responses to stressful events (e.g. Cushing’s syndrome; Sapolsky et al., 2000). Habituation of HPA axis to repeated stressors are also common and this leads to lowering of cortisol levels or blunted diurnal cortisol profile (Thoma et al., 2017). Additionally, higher stress and cortisol level has been found to negatively affect hippocampus which is the major memory control centre (Brown et al., 1996). Therefore, not only does the HPA axis functions in maintaining the basal and stress-related homeostasis but also regulates emotional and cognitive centres in the brain. Thus, for students, chronic stress is likely to interfere with their present academic performance as well as affect long term physical and mental health. However, functioning of basal and reactive responses of the HPA axis is affected by a multitude of other factors from age, sex, dietary intakes, early life experiences, and social factors as well as steroid hormone levels and subjective psychological stress responses (Hsiao et al., 2011). Thus, how students respond to stressors will be highly variable, depending on all the other associated factors that influence their lives.

Psychological stressors are among the most important factor affecting HPA axis activity but extensive research connecting HPA axis reactivity and perceived stress responses have found variable relationships (Halford et al., 2012). In some cases, perceived control over stressful events and perception of stress of an immediate stress stimulus have been shown to affect psychological and physiological responses (e.g. Halford et al., 2012). In other cases, negative or no correlation between physiological responses especially cortisol levels and psychological or subjective stress measures are found (e.g., Halford et al., 2012). Some of this variation can also be attributed to the psychological traits being measured as all parameters are not directly influenced by physiological responses or vice versa. The major psychological measures used across studies are perceived stress scale (PSS), active coping measures, mood scores as well as anxiety-depression scores (Halford et al., 2012). Among these, PSS has been used extensively and it measures the degree to which situations in one’s life were appraised as stressful during the last month. Similarly, positive and negative mood scores also help to quantify the overall mood of an individual over the last month (Watson et al., 1988). Though such psychological measures are an effective way to understand the major causes of acute or chronic stress, the inconsistency in relationship between physiological and psychological responses is not surprising, given the fact that there is a complex neurobiological interplay between perceived stress and HPA axis functioning. Thus, to better understand and assess stressful life events for an individual, both measurements of physiology and psychology will provide a more comprehensive model approach. On one hand perceived stress measures from psychological surveys help quantify the causes of stress and provides an overall idea of chronic stressors over a month-long period whereas immediate responses to acute stressors are captured by cortisol measures which also helps to assess potential health risks in the long-term.

Salivary cortisol has been repeatedly used across studies as a biochemical marker for stress as it can be easily and non-invasively collected. Early morning cortisol levels have been observed to be lower than typical following post-traumatic stress disorder (Wessa et al., 2006), exhaustion (Mommersteeg et al., 2007) and depression (Stetler and Miller, 2005), and elevated responses has been observed in individuals experiencing high work stress (Schulz and Schlotz, 1999). Most studies have been investigating stress and cortisol responses under laboratory conditions by inducing stressful stimuli such as the Trier Social Test or other social challenges (Ellenbogen et al., 2010; Entringer et al., 2010; Espín et al., 2016; Hakamata et al., 2013; Schlotz et al., 2011). Very few studies till date have evaluated stress under real-life conditions and across a longitudinal or repeated scale (Bardi et al., 2011; González-Cabrera et al., 2014).

The present study was conducted on a group of residential undergraduate students of biology majors for an entire year. We selected students during their first year of joining the academic program to understand how students cope with a change in both their academic and personal environment. We measured psychological stress parameters and salivary cortisol in students with repeated sampling of 6 times during the early morning hours before breakfast to understand the relationship between physiological and psychological measures of stress across the repeated sampling events. We hypothesised that stress perception within-individuals would be associated with elevation in cortisol responses. We also anticipated differences in cortisol and perceived stress responses between men and women (similar to other findings by Austin et al., 2018; Dawson et al., 2014). We finally predicted a decrease in perceived as well as physiological stress response along the longitudinal scale as individuals were expected to adjust to the novel academic and social environment over time. To our knowledge this is the first longitudinal study to test both perceived and physiological stress response in undergraduate college students from India.

## Material and methods

### Participants

Twenty-five undergraduate residential students from the biology major participated in this longitudinal study. Participation for this study was voluntary and we had repeated measurements for ~20 individuals across each time point (Men=7, Women=15; age= 17-21years; Time points=6). The study was conducted from August 2018 to May 2019 with sampling done in the months of August (1), September (2), November (3), January (4), March (5) and May (6). We collected 3 samples in the first semester (August-November) and 3 samples in the second semester (January-May). The two sampling points of 5 and 6 at the end of the second semester (March and May) were during assignment submission and during the end-of-the-year examinations.

### Procedure

We measured cortisol through salivary measurements and perceived stress through questionnaires. Participants provided saliva samples before breakfast between 08:00-08:30h and we ensured that individuals did not eat, drink or brushed their teeth 30 mins prior to providing the samples. Participants were requested to passively accumulate and provide ~1-2ml saliva in conical 10ml centrifuge tubes. All samples were stored at −20°C for further analysis. On the same day of saliva collection participants also filled out questionnaires corresponding to perceived stress. The same protocol was used for all repeated measurements. We did not control for menstrual cycle phase for the women participants as early morning cortisol responses are not expected to be significantly affected by menstrual phase (Kudielka and Kirschbaum, 2003).

### Salivary cortisol

Before cortisol analysis, all samples were thawed and centrifuged at 3500rpm for 20 min and the supernatant was used for further analyses. Enzyme-Immuno Assay kits (Arbor Assay DetectX Cortisol K003-H5) were used to measure circulating cortisol level. EIA kits were first optimized (Wada et al., 2007) and we subsequently analysed samples at a dilution ratio of 1:4 in duplicate across 4 assays. Percent recovery of cortisol in the assay was 98.93, with an intra-assay coefficient of variation of 0.12-6.84 and an inter-assay coefficient of variation at 9.51 (Inter-assay CV were calculated from a lab standard of known concentration placed on all plates). Hormone levels were determined in reference to seven-point standard curve with a limit of detection at 0.016 ng/ml for cortisol.

### Psychological evaluation

All participants completed questionnaire on the same day as saliva collection. Three self-reported subjective measures of psychological state were quantified from the questionnaire data: K10 distress scale (Kessler et al., 2002), Perceived Stress Scale-14 (Cohen et al., 1983) and positive mood measure (Watson et al., 1988). K10 was calculated on a 5-point scale with distress or K10 measures having 10 questions on how often individuals felt tired/nervous/distressed with scoring from “all of the time” to “none of the time”. Positive mood score was calculated similarly on a 5-point scale from a total of 13 questions, with individuals scoring how inspired/peaceful/satisfied they felt over the last month on a scale from “extremely” to “none at all”. Perceived stress scale (PSS) comprised of a 14-item questionnaire with scores from 0-4 describing how often individuals felt a certain way in the last month. PSS14 included both positive and negative items and for analysis, the positive scores were reversed before calculating the final PSS score (Cohen et al., 1983). Along with the above three measures we also asked participants one open ended question - what aspect of their life caused maximum stress in the last month: academic, own health, health of close one, relationship stress, family issues, financial issues or any other. General data on health issues and medical history were also obtained. We excluded one individual who was on medication as this would severely affect their cortisol response. Sample of the study questionnaire is provided in the supplementary material.

### Ethical consideration

Participation in this study was voluntary, and the informed consent form was signed by participants at all time points during the sampling. The study was performed in accordance with the Declaration of Helsinki and was approved by the ethics committee of Azim Premji University.

### Statistical Analyses

Individuals were sampled at 6 time points and cortisol, PSS14, K10 and Mood scores were quantified at each. To test whether cortisol, PSS14, K10 and Mood were independently different between sexes across the six different time points, we performed separate linear mixed effect model analyses (R package: lmer and lmer Test, Kuznetsova et al. 2017) for all response variables except the K10 Distress scale where we used generalised linear mixed effect modelling as data was non-normal (R package: glmmADMB, Bolker et al. 2012). We also log transformed cortisol data to normalise it before performing the linear mixed effect models. In all models, sex and time points were used as interacting fixed factors and individual identity as random effect. All individuals did not participate across all time points and thus we avoided using a parametric repeated ANOVA for the analyses and instead used a mixed model framework which would account for differences in sample repeats.

Further to understand if physiological (Cortisol) and psychological (K10, Mood, PSS14) parameters were correlated, we first tested if all psychological parameters were independent or correlated using Pearson’s correlation. We found that all variables were significantly correlated with each other (PSS14 and K10: t=6.12, p<0.001, Pearson’s correlation coefficient=0.48; PSS14 and Mood: t=-6.15, p<0.001, Pearson’s correlation coefficient=-0.48) and thus we only used PSS14 to test for correlation between psychological and physiological responses (cortisol). We used Pearson’s correlation analyses for each time point and sex. We finally scored presence and absence of various types of stressors (based on the open-ended question): academic, own health, health of close one, relationship stress, family issues and financial issues and used a generalised linear mixed effect modelling with a binomial distribution to test which stressor type contributed most across each time point for both sexes. For this, data was divided across sexes and we ran two separate GLMMs, with presence or absence (1/0) as our response, time point and stressor type as our fixed factors and individual identity as random effect. All post-hoc comparisons were performed using lsmeans function (package: lsmeans; Length 2016) and all statistical analyses was performed using R version 3.6 (R core team 2019).

## Results

### Psychological and physiological stress across time

We found sex differences across time points for cortisol response and all other psychological variables except the mood parameter. There was a significant interaction effect of sex and time point for the K10 Distress scale wherein women were not different across all time points (all *t*<2.50, *p*>0.05, Fig. 1a) but men showed decreased levels of distress from time points 1 to 4 (*t*=3.86, *p*=0.002, Fig. 1a), 1 to 6 (*t*=2.94, *p*=0.044, Fig 1a) and also 2 to 6 (*t*=2.98, *p*=0.039, Fig 1a). Similarly, for PSS14 we found no difference across time points for women (all *t*<1.85, *p*>0.05, Fig. 1b) but men showed a decrease in PSS14 from time points 1 to 5 (*t*=2.96, *p*=0.046, Fig. 1b) and 2 to 5 (*t*=2.98, *p*=0.041, Fig. 1b). There was no significant difference between the sexes or across time points for Mood scores (Fig. 1c). Thus, we found that men, but not women showed a reduction in perceived level of stress (PSS14) and distress (K10) across time.

**Figure 1.**
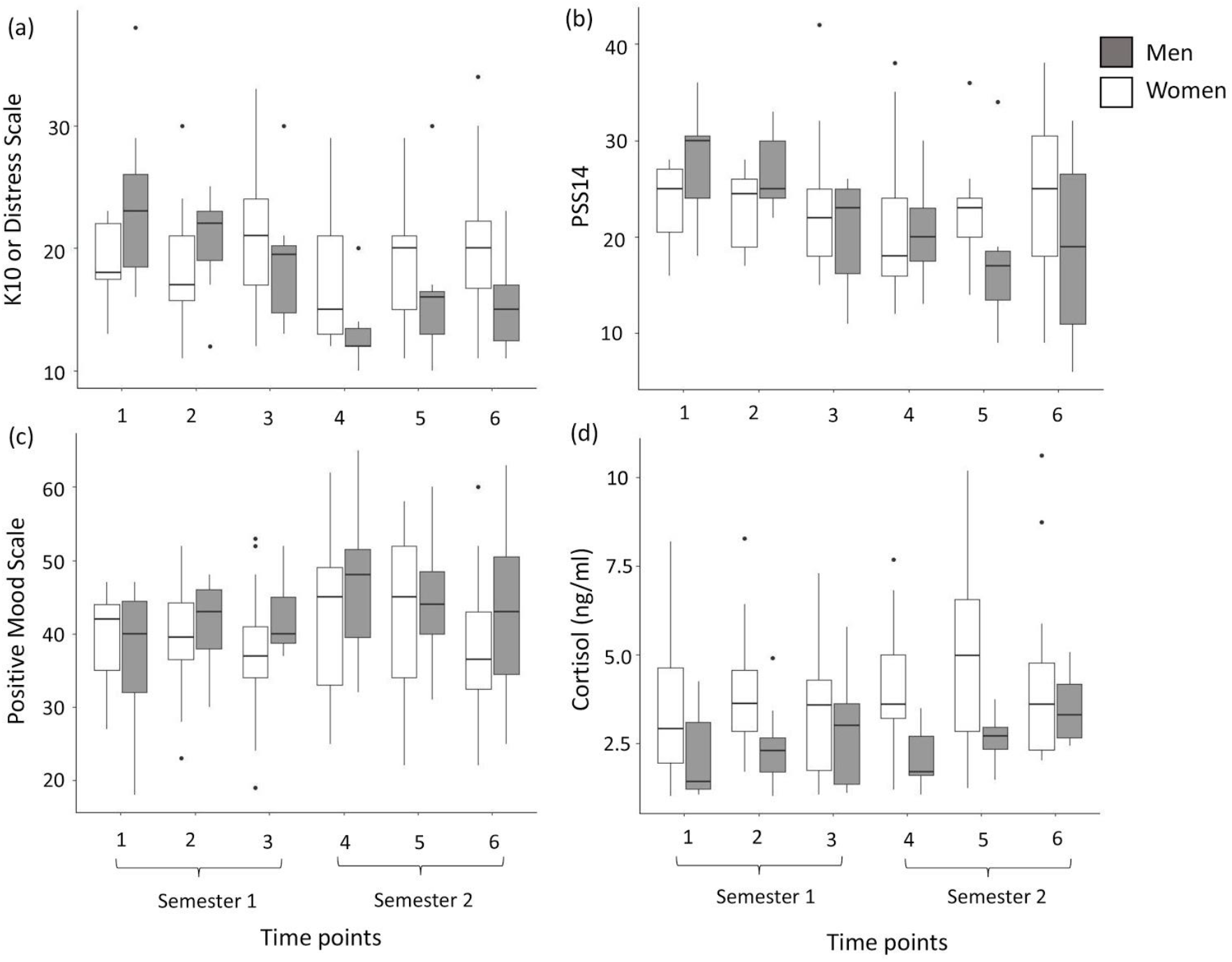
Psychological measures of (a) K10 Distress scale, (b) PSS14, (c) Mood scale and physiological measures of (d) Cortisol levels of men and women across all time points. Grey and white boxes represent responses of men and women respectively. Boxplots show medians, quartiles, 5^th^ and 95^th^ percentiles and extreme values.

Similar to the psychological stress responses, there was a significant difference between men and women in cortisol response with men having overall lower levels of circulating cortisol compared to women (*t*=-2.05, *p*=0.042, Fig. 1d). We also found cortisol levels to be significantly higher for women at time point 5 compared to the first time point (*t*=2.22, *p*=0.028, Fig. 1d).

Contribution of individual identity or random effect across all models was 1.5 standard deviation or lower.

### Correlation between physiological and psychological stress responses

When tested separately across sexes and across all time points, we found no significant correlation between PSS14 and cortisol for any combination of sex and time (all p>0.05).

### Types of stressors

All students reported binary (yes/no) responses for presence or absence of different types of stressors which were academic, own health, health of close ones, relationship, financial and family. There were no sex differences across types of stressors reported (z=-0.66, p=0.509), but different type of stressors significantly differed across time. We thus divided the data across sexes and performed separate mixed effect models to understand how the stressors were different across time points. Total number of stressors reported by both men and women were lowest at time point 4 compared to time point 1 (men: z=-2.5, p=0.012, Fig. 2a; women: z=-2.86, p=0.004, Fig. 2b). Additionally, men also reported a significantly lower number of stressors at time point 5 compared to 1 (z=-2.23, p=0.025, Fig. 2a).

**Figure 2.**
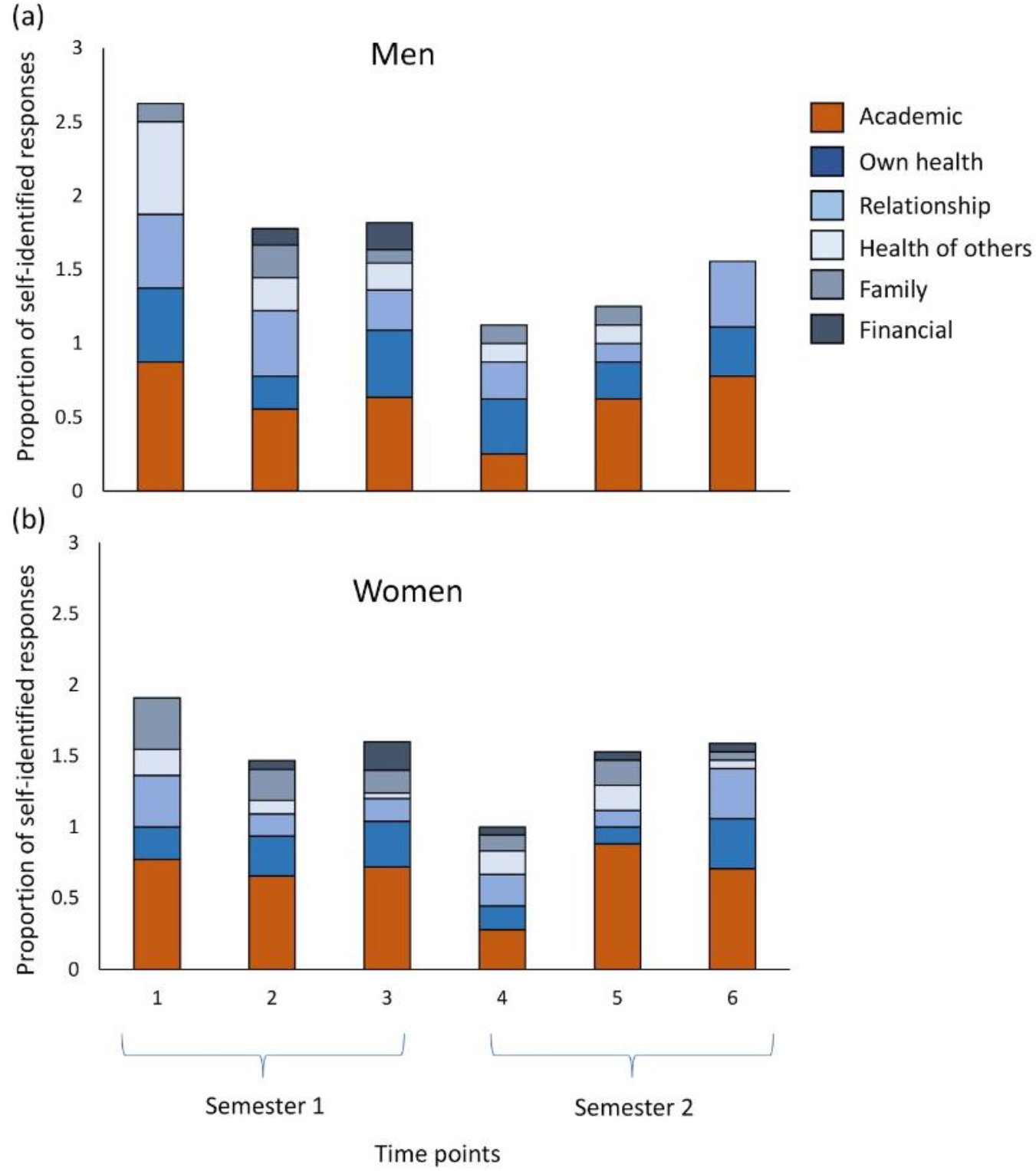
Stacked bar plot shows proportion of self-identified responses for the different types of stressors reported across all time points by (a) men and (b) women participants.

Academic stressor was reported by both males and females the greatest number of times compared to all other stressor types (all z>3, p<0.03, Fig.2). After academic stressor, own health was reported to be next highest compared to financial stress which was reported least number of times (men: z=3.02, p=0.026, Fig. 2a; women: z=3.04, p=0.024, Fig. 2b). Men also reported relationship stress more often than financial stress (z=2.90, p=0.037, Fig. 2a).

## Discussion

The current study provides a combination of measures from both psychological or perceived stress as well as physiological or cortisol responses of residential undergraduate students across a yearlong period. Both men and women students participated in the study and we found that overall, men had lower salivary cortisol levels across the year compared to women. Men also reported a lower score of perceived stress and distress as time progressed after joining the first year of undergraduate education. Women participants reported similar levels of perceived stress and distress across the year although their cortisol responses increased during the end of the semester when assignment submissions were due. When asked about the different types of stressors, students reported academic stress to be the most prevalent as this was reported the maximum number of times across all time points. Financial stress was reported the least number of times. Notably, we found no direct correlation between psychological stress perception and physiological stress responses (cortisol).

In humans, the end product of the stress-responsive neuroendocrine system or HPA axis is cortisol. Cortisol acts as the major regulatory hormone that mediates resource allocation during any stressful condition. Salivary cortisol is unbound by glycoproteins and therefore biologically active, and thus provides a good measure of the stress reaction of an individual (Fink, 2000; Tsigos and Chrousos, 2002). Stressful stimuli can activate HPA axis functioning, which in turn elevates cortisol levels (McEwen, 1998). Interestingly, most of the human psychological stress studies have either found no significant difference between the sexes or have found that young men have elevated cortisol levels compared to young women when challenged with acute stressful tasks such as examinations or other laboratory stress tests (see review: Kudielka and Kirschbaum, 2005). This pattern is contrary to our results where we find that women have higher salivary cortisol levels compared to men across an entire year. However, unlike most previous studies, our study does not specifically induce any stressor to the participants and the salivary cortisol represents an unstimulated daily level. Two of the sampling points in our study were during assignment submission stages and we find an increase in salivary cortisol during one such time point (time 5, Fig 1d) compared to the initial time when there was no immediate academic pressure. This elevation in cortisol is likely to be attributed to the anticipation of stressful events (assignment deadlines and marks). The level of elevated salivary cortisol found during this time point in our study is similar to other studies where students are found to increase to comparable cortisol levels during examination (González-Cabrera et al., 2014; Singh et al., 2012). Stressful stimuli or stressful environments trigger both physiological and psychological responses and because both of these are indicators and outcomes of the same phenomenon, we expected some level of correlation between the two. However, we found no correlation between cortisol and other psychological variables, which is similar to the lack of association found in other studies (see reviews: Halford et al., 2008). This can be majorly attributed to the fact that salivary cortisol levels capture the current state whereas PSS14 or the K10 Distress scale used in our study captures perceived stress over a month-long period. Similar results are also observed in a large number of other studies which use PSS14 or PSS10 as a measure for psychological stress (Dawson et al., 2014; Manigault et al., 2018; Putterman and Linden, 2006), whereas the studies using immediate perceived stress measures such as Visual Analog Scale or Stress-O-meter tend to find a correlation between salivary cortisol and perceived stress measures (Chellew et al., 2015; Chong et al., 2017; Esch et al., 2007; Linnemann et al., 2015; Myint et al., 2011). Men in our study showed a decrease in both perceived stress score (PSS14) and distress score (K10) with progression of time. During semester 2, men reported significantly lower scores for perceived stress whereas women did not change their perception of stress from the start of the semester to the end of the academic year. This suggests that men might be adjusting to the new academic environment faster than women. Irrespective of the generally lower perception of stress by women and men, cortisol levels were highest during assignment submission and the end-of-the-year-examination time compared to the start of the semester. This further strengthens the view that cortisol level is majorly influenced by immediate stressors compared to perceived stress measures which are representation of a much longer time period.

The major contributing factor for perceived stress in our study seems to be academic pressure as that was reported most often by both men and women across all time points. Overall stressors reported was lowest at the beginning of semester 2 when academic pressure was low and students returned to classes after a break. Though a similar pattern was expected at the beginning of semester 1, the anticipation of a new academic environment and the shift from high school to a college environment likely led to a heightened perception of academic stress at the beginning of semester 1. In some previous studies, financial stress has been reported to be a cause of major stressor for undergraduates coming from different socio-economic backgrounds (Kumar et al., 2009; Morra et al., 2008). However, in our study, we find financial stress to be reported least number of times and thus of least concern, which is most likely due to the financial security provided by the University through scholarships. Further, since students stay within a residential campus, they are partially protected from the daily exposure of individual and potentially, familial financial stress. It would be interesting to compare stress profiles of students studying under similar environment but from residential and non-residential study programs.

One major limitation of our study is that we did not have information on smoking behaviour or any other consumption of drugs or alcohol. While these variables can influence physiological responses, our repeated measures design should ameliorate the influence of such variables on our findings. We also did not include any questions on the menstrual cycle phase for women because previous studies report that morning cortisol responses were not altered by menstrual cycle phase (Kudielka and Kirschbaum, 2003). However, there have been other studies which also report that women in their luteal phase have similar cortisol responses as men, but women who are in their follicular phase or taking oral contraceptives tend to have lower cortisol responses compared to men (Clemens et al., 1999). In the current study, menstrual phase is unlikely to have influenced our findings and we find that women had higher cortisol responses than men across all time points despite the lack of including menstrual phase information as a covariate. We intentionally excluded information on diet, as our study was on residential campus, and thus all college students were provided the same food during the entire study duration. Finally, we had a sex-biased sample where there were more women than men in our study group and although we had a longitudinal study design, we obtained single measurements at each time point. Future studies should ideally have multiple measurements at each time point with a sex-balanced design along a longitudinal scale. Despite these potential weaknesses, the importance of measuring both physiological and psychological measures are clear, as we observe different patterns of cortisol and perceived stress across different time points in the academic semester. Future research using multiple variables for stress measurements with a greater sample size and longitudinal design would help in addressing the growing issue of poor mental health in academia. With such information on stress-induced triggers and when to expect them, we can also design optimal intervention strategies for a healthy young adult population.

The findings of the current study show that among first year undergraduate students from an Indian University, men experience lower perceived stress with time spent in a new academic environment and also exhibit significantly lower cortisol responses compared to women. Academic stressor was perceived as most significant and financial as least significant stressor in the first year of college.

## Acknowledgements

The authors would like to thank the 2018 BSc Biology students of Azim Premji University for volunteering for this study. We would also like to thank Anagha Menon and Elizabeth Matthew for help with collecting saliva samples. We would also like to thank Azim Premji University for funding this study.

