## Supplementary material for "A longitudinal study of perceived stress and cortisol responses in an undergraduate student population from India"

PLEASE ANSWER AS ACCURATELY AS POSSIBLE. PLEASE REMEMBER THAT YOUR RESPONSES WILL HAVE AN IMPACT ON THE OVERALL STUDY FINDINGS ON COLLEGE YOUTH.

THANK YOU…..

**Name**: **Age**:

**Gender**: Male/Female **Religion:** Hindu/ Muslim/ Christian/ Sikh/ others (specify)

**Course name & year**:

**Any chronic medical condition: Yes/ No, If yes, please specify.**

**Currently seeking any professional help for any Medical problem: Yes/ No/ Not Applicable**

**Do you feel that you are currently suffering from any psychological problem? (E.g. anxiety, depression)** No/ Yes (please specify)

**Have you participated in any program/courses on personal growth/ self development at any point of time?** No/ Yes

If Yes when? Please specify the nature of the program:

**Have you consulted a health professional for any kind of psychological problem at any point of time?**

Yes (any time in the past before 6 months) Yes (currently in the last 6 months) No

**Have you taken medication for PCOD/PCOS? (Polycystic ovarian syndrome)**

Yes (can you please provide details about when and which medication) No

**Would you want or prefer to be involved in some activities which can help in enhancing growth and functioning of students? Yes / No / Do not know**

**If yes, please mention what you will be interested in?**

**In the past 4 weeks about how often did you feel…**

**(**Please tick the box that most closely fits)

| **K10** | | All of the time | Most of the time | Some of the time | A little of the time | None of the time |
|---|---|---|---|---|---|---|
| 1 | tired out for no good reason? | | | | | |
| 2 | nervous? | | | | | |
| 3 | so nervous that nothing could calm you down? | | | | | |
| 4 | hopeless? | | | | | |
| 5 | restless or fidgety? | | | | | |
| 6 | so restless you could not sit still? | | | | | |
| 7 | depressed? | | | | | |
| 8 | that everything was an effort? | | | | | |
| 9 | so sad that nothing could cheer you up? | | | | | |
| 10 | worthless? | | | | | |

**My mood in the PAST FEW WEEKS**

| | | Extremely | Quite a bit | Moderately | A little | Not at all |
|---|---|---|---|---|---|---|
| 1 | Satisfied and content | | | | | |
| 2 | Pleasantly at rest | | | | | |
| 3 | (feeling) excited | | | | | |
| 4 | Peaceful | | | | | |
| 5 | Very inspired | | | | | |
| 6 | Calm and relaxed | | | | | |
| 7 | Powerful and strong | | | | | |
| 8 | Set and determined about something | | | | | |
| 9 | Sharp and attentive | | | | | |
| 10 | (feeling) enthusiastic | | | | | |
| 11 | At ease and comfortable with things | | | | | |
| 12 | Alive and active | | | | | |
| 13 | Proud of myself | | | | | |

**<u>Sources of stress in the last 3 months</u>**

**Please place a tick mark against stressors that have been most troublesome for you in the last 2 months and give a short description:**

1. **Related to academics**

2. **Related to my own health**

3. **Related to health/ illness of a person close to me**

4. **Conflict/ strain in close relationship**

5. **Financial issues**

6. **Family issues**

7. **Any other**

### My level of Stress in the last one month

**The questions in this scale ask you about your feelings and thoughts during THE LAST MONTH. In each case, you will be asked to indicate your response by placing an "X" over the circle representing HOW OFTEN you felt or thought a certain way. Although some of the questions are similar, there are differences between them and you should treat each one as a separate question. The best approach is to answer fairly quickly. That is, don't try to count up the number of times you felt a particular way, but rather indicate the alternative that seems like a reasonable estimate.**

| | Never<br>1 | Almost Never<br>2 | Sometimes<br>3 | Fairly Often<br>4 | Very Often<br>5 |
|---|---|---|---|---|---|
| **1. In the last month, how often have you been upset because of something that happened unexpectedly?** | ○ | ○ | ○ | ○ | ○ |
| **2. In the last month, how often have you felt that you were unable to control the important things in your life?** | ○ | ○ | ○ | ○ | ○ |
| **3. In the last month, how often have you felt nervous and "stressed"?** | ○ | ○ | ○ | ○ | ○ |
| **4. In the last month, how often have you dealt successfully with day to day problems and annoyances?** | ○ | ○ | ○ | ○ | ○ |
| **5. In the last month, how often have you felt that you were effectively coping with important changes that were occurring in your life?** | ○ | ○ | ○ | ○ | ○ |
| **6. In the last month, how often have you felt confident about your ability to handle your personal problems?** | ○ | ○ | ○ | ○ | ○ |
| **7. In the last month, how often have you felt that things were going your way?** | ○ | ○ | ○ | ○ | ○ |
| **8. In the last month, how often have you found that you could not cope with all the things that you had to do?** | ○ | ○ | ○ | ○ | ○ |
| **9. In the last month, how often have you been able to control irritations in your life?** | ○ | ○ | ○ | ○ | ○ |

| | Never<br>1 | Almost Never<br>2 | Sometimes<br>3 | Fairly Often<br>4 | Very Often<br>5 |
|---|---|---|---|---|---|
| **10. In the last month, how often have you felt that you were on top of things?** | ○ | ○ | ○ | ○ | ○ |
| **11. In the last month, how often have you been angered because of things that happened that were outside of your control?** | ○ | ○ | ○ | ○ | ○ |
| **12. In the last month, how often have you found yourself thinking about things that you have to accomplish?** | ○ | ○ | ○ | ○ | ○ |
| **13. In the last month, how often have you been able to control the way you spend your time?** | ○ | ○ | ○ | ○ | ○ |
| **14. In the last month, how often have you felt difficulties were piling up so high that you could not overcome them?** | ○ | ○ | ○ | ○ | ○ |
